# Rapid Invisible Frequency Tagging (RIFT) with a consumer monitor: A proof-of-concept

**DOI:** 10.1101/2025.08.14.670287

**Authors:** Olaf Dimigen, Ioana Badea, Iarina Simon, Mark M. Span

## Abstract

Rapid Invisible Frequency Tagging (RIFT) enables neural frequency tagging at rates above the flicker fusion threshold, eliciting steady-state responses to flicker that is almost imperceptible. While RIFT has proven valuable for studying visuospatial attention, it has so far relied on costly projector systems, typically in combination with magnetoencephalography (MEG). The recent emergence of high-speed organic light-emitting diode (OLED) monitors for consumers suggests that RIFT may also be feasible with much more accessible hardware. Here, we provide a proof-of-concept demonstrating successful RIFT using a consumer-grade 480 Hz OLED monitor in combination with electroencephalography (EEG). We also share practical recommendations for achieving precise stimulus timing at 480 Hz with minimal frame drops. In a central fixation task, participants viewed a tapered disc stimulus flickering either centrally or peripherally. Luminance was modulated sinusoidally at 60 Hz or 64 Hz, frequencies at which the flicker was barely visible. Photodiode recordings confirmed that the monitor delivered accurate frame timing with few dropped frames. Cross-coherence analysis between occipital EEG channels and a photodiode revealed robust, frequency-specific neural tagging responses for central stimuli at both frequencies. In comparison, weaker coherence was observed for 60 Hz peripheral flicker. Our findings demonstrate that RIFT can be reliably implemented using affordable stimulation hardware, a low-density EEG montage, and a minimal processing pipeline. We hope that this lowers barriers to entry, facilitating broader use of RIFT in basic research and in applied settings where cost and portability matter.

---

Rapid invisible frequency tagging (RIFT) is a technique that uses high-frequency flicker to elicit neural steady-state responses in the visual cortex, while avoiding the perceptually salient stimuli typical of traditional steady-state visually evoked potentials (SSVEPs; Regan, 1977). SSVEPs have long been used to track visuospatial attention using magneto- or electroencephalography (M/EEG). However, unlike most SSVEP paradigms that rely on slower, binary flicker in the range of 5–20 Hz, RIFT smoothly modulates stimulus luminance or contrast at frequencies of 60-84 Hz (Minarik et al., 2023). Because these frequencies are above typical fusion thresholds, the flicker remains barely visible or even imperceptible (Spaak et al., 2024) to the participant. This minimizes distraction and fatigue, making the technique attractive for the continuous tracking of visuospatial attention under more naturalistic conditions (e.g., Arora et al., 2025, Pan et al., 2021; Zhigalov et al., 2019; for review see Seijdel et al., 2023).

Despite these advantages, the widespread adoption of RIFT has been limited by technical and financial constraints. With a single exception (Arora et al. 2024; see also Arora et al., 2025), all published RIFT studies to date have used MEG for neural recordings. Furthermore, as mentioned, the low visibility of the driving stimulus is partly achieved by modulating its luminance or contrast in a sinusoidal fashion rather than toggling it on and off. Displaying this smooth time series has required state-of-the-art DLP LED projectors (i.e., the PROPixx by VPixx Technologies) with refresh rates of up to 480 Hz (color) or 1440 Hz (greyscale). While these dedicated systems offer precise frame timing, they cost over €20,000 and require setups only available in well-funded MEG or vision laboratories. Thus, even in recent studies using EEG (Arora et al. 2024; 2025), stimulation still relied on a specialized projector, leaving a major hardware bottleneck unresolved.

A promising avenue for democratizing RIFT lies in recent improvements in consumer display technology. Specifically, organic light-emitting diode monitors (OLEDs), which are now available with refresh rates of up to 500 Hz, offer key advantages over flatscreen monitors of the past (Dimigen & Stein, 2024). For example, due to their self-emissive pixels, OLEDs have rapid pixel response times (<0.3 ms) and can show true blacks. This allows them to produce high-contrast luminance changes with minimal temporal smearing, making them a suitable candidate for RIFT (Dimigen & Stein, 2024).

Crucially, the cost of these high-speed OLEDs, developed for gaming, is now below 1,000 EUR, an order of magnitude lower than that of specialized projectors. By dramatically lowering the hardware barrier, OLEDs in combination with EEG have the potential to make RIFT more widely available and extend the approach also to brain-computer interfaces (BCI; Brickwedde et al., 2022; see also Ladouce et al., 2022) or clinical applications where cost and portability are key.

In this pilot study, we tested the feasibility of RIFT with off-the-shelf display hardware. Specifically, we asked whether a 480 Hz OLED, in combination with a low-density EEG montage and a minimal preprocessing pipeline, is sufficient to capture frequency-specific neural tagging responses to barely visible flicker. Our goals were twofold: (1) To document a working OLED-based RIFT setup with reliable timing, and (2) to evaluate whether this setup elicits reliable neural tagging responses in EEG.

## METHODS

### Participants

Ten young adults (20-22 years old, M = 21.2, SD = 0.632, 7 female), with normal uncorrected vision, were paid 20 € for participation. All participants reported no neurological or psychiatric illness, including no history of photo epilepsy. The study was approved by the ethics committee of the Behavioural and Social Science faculty of the University of Groningen (request PSY-2425-S-0321) and participants provided written informed consent.

### Stimulus

Stimuli were presented on the ASUS PG27AQDP (firmware: MCM103) OLED, running at 480 Hz and a 1920 × 1080 pixel resolution. The monitor was gamma-calibrated (linearized) to a Gamma of 1 using a Konica-Minolta LS-150 photometer. Stimuli were presented using Psychtoolbox (v.3.0.19.16) under MATLAB (R2024b). The monitor was allowed to warm up for 30 min before recordings (Dimigen & Stein, 2024).

As the RIFT stimulus, we used a luminance-modulated disc (circular patch) with a soft edge, which oscillated between 0% black (RGB-level: 0; luminance: 0 cd/m^2^) and 100% white (RGB-level: 255; 219 cd/m^2^) during each cycle. The disc therefore appeared on average as gray and invisible against the gray (50%) background (RGB-level: 128; 109 cd/m^2^). Ambient illuminance in the recording room was 80 lux. At the 55 cm viewing distance, maintained with a chin rest, the disc’s central full-contrast region extended to a radius of r=2.55°, whereas its soft edge, tapered with a cosine-shaped alpha taper, fully faded into the background at r=7.34°. A wide taper was used to decrease flicker visibility in the presence of (fixational) eye movements (see Discussion).

In the *central* condition (used with 60 Hz and 64 Hz flicker; see also Arora et al., 2024), the disc was centered on the fixation dot. In the *peripheral* condition (only explored here with 60 Hz flicker and in a subset of 7 participants^1^), the disc’s center was shifted 12° to the left. This meant that the inner (right) edge of the disc’s full-contrast region was located at 9.43° eccentricity.

Across time, the disc’s luminance was modulated in a (quasi) sinusoidal manner. In most prior studies, 60 Hz RIFT was discretized into 24 frames/cycle at 1440 Hz. With the 480 Hz OELD, we could show 60 Hz flicker as 8 frames/cycle. The 64 Hz flicker does not divide into an integer number of frames, but to 7.5 frames/cycle. To solve this, two cycles of a 64 Hz sinusoid were mapped onto 15 subsequent frames (cf., Arora et al., 2024; note: this produces a subharmonic at 32 Hz, but this should not be a problem for most RIFT applications).

### Procedure

The experiment comprised 99 trials belonging to the three conditions (33 trials each). Each trial lasted about 10 seconds, during which a central black fixation dot (5 × 5 pixels) was continuously presented in the screen center. Participants were asked to maintain a steady fixation on the dot, refrain from blinking if possible, and to blink during self-paced breaks after each trial (terminated by a keyboard press).

During each trial, a 60 Hz or 64 Hz stimulus was shown for 600 cycles (10 s at 60 Hz; 9.38 s at 64 Hz), with the central 500 cycles presented at full contrast. To reduce onset transients and visibility, the stimulus faded in and out over 50 cycles (883 ms at 60 Hz), using a raised cosine taper applied multiplicatively to the sinusoidal luminance modulation. Examples are shown in Figure 1C.

**Figure 1.**
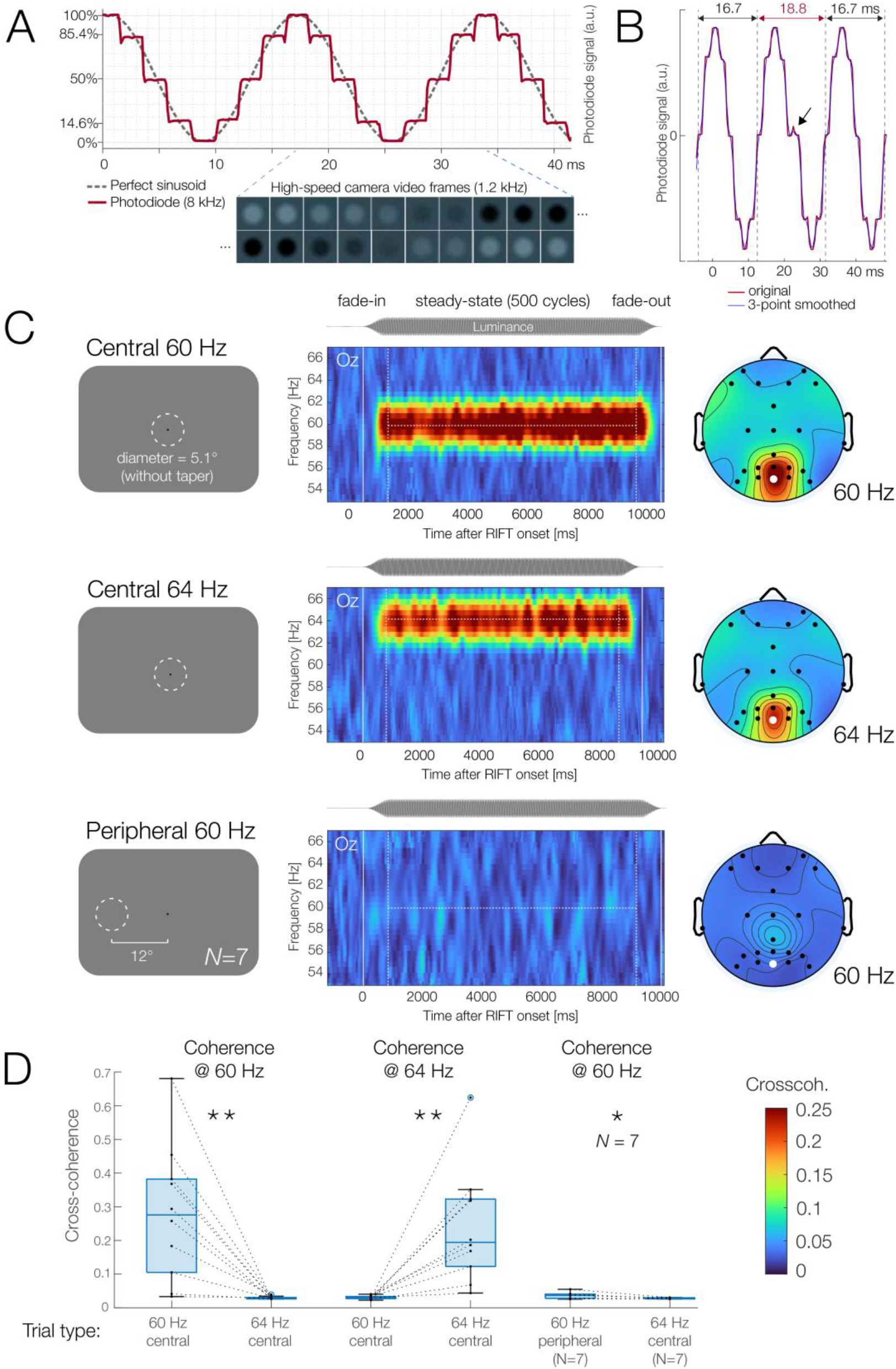
(A) RIFT luminance time series (here: 60 Hz at 8 frames/cycle) displayed on the 480 Hz OLED, as measured with a photodiode and high-speed camera. The monitor rapidly and accurately reached each luminance state. (B) Rare example of a dropped frame (black arrow), detected via a longer interval between rising zero-crossings (vertical dotted lines). (C) Neural tagging results. Topographies show mean cross-coherence during the steady-state interval. Oz is highlighted in white. (D) Statistics confirm significant RIFT responses in all three conditions.

Parallel port triggers were sent to the EEG at multiple moments during each trial, including at onset of the fade-in. To track the luminance output, a BrainProducts *PhotoSensor* photodiode was attached to the upper left screen corner which displayed a copy of the RIFT stimulus and its signal was fed into the EEG amplifier’s external inputs. Timestamps for each frame (e.g., VBLtime) were also logged by Psychtoolbox.

The 99 trials were shown in an individually randomized order. After the experiment, participants were asked whether or not they had perceived any flicker.

### Monitor & System configuration

We tested numerous configurations of the stimulation computer, operating system, and monitor settings before finding a suitable setup that ensured accurate stimulation at 480 Hz with few dropped frames. Below we briefly report the feasible settings for researchers interested in replicating our setup.

Stimuli were presented on a PC running Ubuntu Linux (version 24.04.2 LTS; kernel: v.6.11.0-26-generic) equipped with an AMD RX6600 GPU, an Intel Core i7-12700 CPU, and 32 GB of RAM. The Ubuntu system ran MESA drivers (v24.0.7) using GNOME in an X.Org session (v21.1.11) with Wayland deactivated^2^. The monitor was connected via Display Port (v.1.4) and operated at a “Brightness” setting of 80%, a “Contrast” setting of 80%, a color depth of 18 bit, and with the “Racing Mode” presetting. Variable refresh rate (VRR) and High Dynamic Range (HDR) were deactivated. The “Uniform Brightness” mode was activated to avoid Auto-Brightness Limiting (Dimigen & Stein, 2024).

Importantly, we found that it is essential to deactivate digital stream compression (DSC) on the monitor. DSC is a lossless compression used to transmit high-bandwidth data over the video interface; however, we found that it leads to unreliable frame timing at 480 Hz. Deactivation of DSC fixed this issue but also forced the use of a lower full HD resolution (compared to the monitor’s native 2560 × 1440).

### Electrophysiological recording

EEG and electro-oculogram (EOG) signals were recorded from a total of 23 electrodes, mostly placed in an ANT Waveguard cap at standard 10-10 system positions. External electrodes were placed on the infraorbital ridge (IO1, IO2) and canthus (LO1, LO2) of each eye and the mastoids (M1, M2). An additional electrode at AFz served as ground. Signals were amplified using a TSMi Refa (model: Refa8-32e4b4a) DC amplifier, digitized at 2 kHz and recorded against an online average reference. Impedances were kept <10 KΩ using ECI gel.

### EEG preprocessing

Analyses were conducted with custom Matlab scripts using EEGLAB functions (Delorme & Makeig, 2004; v.2024.2). EEG preprocessing was deliberately kept minimal: Offline, the average-referenced data was down sampled to 1000 Hz and high-pass filtered at a -6 dB cutoff of 1 Hz (transition bandwidth: 2 Hz) using EEGLAB’s default FIR filter (*pop_eegfiltnew*.*m*). Aggressive high-pass filtering was applied as it improves ICA decompositions (Dimigen, 2020). Data were then epoched from -2 to +11 s relative to the fade-in onset trigger. ICA was trained on these epochs (*k* = 2,434 data points per weight), and components were auto-classified using *ICLabel* (Pion-Tonachini et al., 2019). Components labeled as “Eye” or “Muscle” artifacts with ≥90% probability were removed (mean = 3.1; range: 2–4). After artifact correction, EOG electrodes (IO1/2, LO1/2) were treated like any other EEG channel. No baseline correction of any type was applied.

### Analysis of stimulation fidelity (dropped frames)

Faithfully updating a stimulus every 2.08 ms without dropping frames (e.g., displaying a frame for longer than one refresh cycle) is challenging on modern graphics hardware, and much more challenging than at 240 Hz (Dimigen & Stein, 2024). To illustrate how precisely modern OLED monitors can track the intended luminance, we also test-recorded 60 Hz flicker with the photodiode sampling at 8 kHz and a Casio EXILIM EF-X1 camera filming at 1200 Hz. Furthermore, the actual experimental data was analyzed for mistimed frames. For this, we focused on the 8.3 s periods of full-amplitude steady-state flicker in the central 60 Hz condition. After smoothing the photodiode signal with a 3-point moving average, dropped frames were detected as outliers (threshold: <16 ms or >17 ms) in the interval between successive rising zero-crossings of the sinusoid, which should last 16.7 ms (1/60 s).

### RIFT analysis: Cross-coherence

Previous studies have used alternative methods to quantify the tagging response. A straightforward approach is to compute the cross-coherence between the EEG and the actual driving video signal captured by the photodiode (magnitude-squared coherence, e.g., Minarik et al., 2023).

We performed a time–frequency decomposition of the epoched data in EEGLAB using a short-time Fourier transform with a 510 ms window and a zero-padding ratio of 4, yielding complex-valued coefficients for each time point and for frequencies between 52 and 68 Hz in

∼0.49 Hz steps. Cross-coherence was then computed between each EEG channel and the photodiode signal using EEGLAB’s *newcrossf*() function with the ‘coher’ option. In this mode, the EEGLAB function returns the magnitude coherence (not yet squared), that is, the absolute value of the complex coherence obtained by averaging trial-wise complex cross-spectra and normalizing by the trial-averaged autospectra. We squared these values to obtain the magnitude-squared coherence (e.g., Minarik et al., 2023), which estimates the proportion of signal power at a given frequency that is linearly shared between the photodiode and the EEG channel. Coherence was computed per channel and condition by averaging spectra across trials; trials were not concatenated.

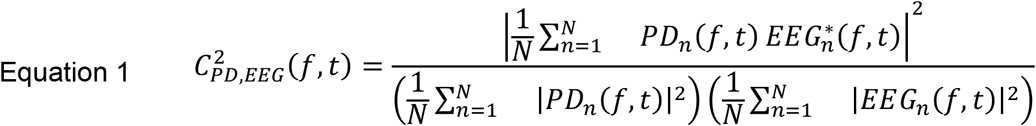

where *N* is the number of trials, *EEGn(f,t)* and *PDn(f,t)* are the complex time-frequency coefficients for the EEG channel and the photodiode (PD) for trial *n*; and ∗ denotes the complex conjugate. The numerator averages the trial-wise cross-spectra before taking the magnitude; the denominator is the product of the trial-averaged autospectra.

### Statistics

For statistical comparisons (Figure 1D), we focused on mid-central electrode Oz, selected a-priori as the classic electrode to capture (SS)VEPs, at least for foveal stimuli. For this electrode, cross-coherence values were averaged across the 500-cycle steady-state period of each trial. Cross-coherence at the actual target frequency (60 Hz coherence in 60 Hz stimulation trials; 64 Hz in 64 Hz stimulation trials) was then compared to cross-coherence at the same frequency but in trials with the *other* stimulation frequency (e.g., 60 Hz coherence in 64 Hz trials or vice versa). For simplicity, we report below statistics based on one-tailed paired t-tests, testing the directed hypothesis that coherence at the target frequency is larger in stimulation trials. Given our small sample, we also repeated the analyses with a non-parametric permutation test based on 5,000 random permutations of 60/64 Hz condition labels. Since highly similar results were obtained with permutations, we only report t-tests below.

## RESULTS

### Flicker visibility

Following the experiment, all participants reported that they barely noticed the flicker and that they did not perceive it continuously; furthermore, four of the participants could not tell at how many screen locations flicker was shown. However, no participant remained entirely unaware of any screen changes throughout the experiment.

### Stimulation fidelity (dropped frames)

Among the more than 1.3 million video frames presented to all participants in the analyzed central 60 Hz condition, we detected 42 mistimed frames (mean per participant: 4.2, range: 0-14, SD: 4.18). In almost every case, a given gray level was presented for two frames instead of one (Figure 1B). At the level of trials, M=10% of our long trials (3.3 trials of 33) contained a dropped frame (range: 0-8. SD: 2.67). In informal tests, single dropped frames did not cause a perceivable visual transient.

### RIFT response

Figure 1C summarizes the neural tagging responses. With central stimulation, we observed clear RIFT responses at both 60 Hz and 64 Hz in the respective stimulation trials, as compared to trials in which stimulation happened at the other frequency. Specifically, at electrode Oz, 60 Hz cross-coherence was larger in central 60 Hz trials (M = 0.280, SD = 0.202) than central 64 Hz trials (M = 0.030, SD = 0.004; *t*(9) = 3.906, *p* < 0.0018, Cohen’s *d* = 1.235). Conversely, 64 Hz cross-coherence was larger in central 64 Hz trials (M = 0.241, SD = 0.171) than central 60 Hz stimulation trials (M = 0.031, SD = 0.005, *t*(9) = 3.833, *p* < 0.0020, *d* = 1.212). Figure 1C also shows results for the 60 Hz peripheral stimulation, presented to a subset of seven participants. Cross-coherence in peripheral trials was generally much weaker (M = 0.036, SD = 0.010), however, this was still a significant increase compared to 60 Hz coherence in the central 64 Hz stimulation trials used for comparison, M = 0.028, SD = 0.002; *t*(6) = 2.430, *p* < 0.0256, *d* = 0.919.

## DISCUSSION

Our results demonstrate that RIFT is feasible with a consumer-grade OLED monitor. Despite our modest sample size, we observed robust and frequency-specific EEG tagging responses to centrally presented, barely visible flicker at 60 and 64 Hz, highlighting the potential of this approach for accessible and cost-effective neural tracking.

Key requirements for implementing RIFT with a consumer monitor are rapid display response times and the accurate timing of frames. At 480 Hz, stimulus updates must happen reliably every 2.08 ms. Here we show that with an appropriate configuration of hard- and software (including deactivating DSC, reducing the resolution, and correcting a Linux bug), stimulus timing problems are rare and limited to the rare dropping of single frames (Figure 1B), which do not seem to affect visibility. Frame drops occur at very low rates (less than 1 in 32,000 frames), enabling excellent stimulation fidelity.

While the setup produced robust responses for central stimuli, responses were markedly weaker albeit still significant for 60 Hz flicker in the periphery (12° eccentricity) shown to a subset of participants. This is replicating prior work showing that coherence strongly decreases with increasing eccentricity (Minarik et al., 2023). Furthermore, it has been shown that RIFT works better for stimuli in the lower visual field (LVF; Minarik et al., 2023), consistent with decades of EEG work showing a LVF-advantage in VEPs (e.g., Capilla et al., 2016), because the LVF projects to more dorsal and upward-facing regions of striate and early extrastriate cortex. To replicate the conditions of many classic attentional cueing studies (e.g., Posner et al., 1980), we chose here to present the peripheral stimulus on the horizontal midline. Moving it to the LVF would have likely enhanced the effect.

Lastly, while all participants described the flicker as barely visible and non-distracting, none remained entirely unaware of it. Psychophysical work on RIFT’s (in)visibility is surprisingly scarce, but Spaak et al. (2024) recently reported complete invisibility of parafoveal (less than about 5° eccentricity) gratings containing sharp edges. In our informal testing with the OLED, we observe that flicker visibility is as expected higher for locations in the periphery, which have a higher fusion frequency (Tyler & Hamer, 1990). We also find that RIFT stimuli with sharp edges that are virtually imperceptible during fixation become highly visible during medium-sized saccades (of about 5°) when the flicker is smeared across the retina at high velocities (cf. Schweitzer et al., 2019). However, strong edge tapering and a strict fixation instruction mitigated this problem.

Spaak et al.’s results of total RIFT invisibility may reflect the higher frame rate of their projector compared to the OLED (24 vs. 8 frames/cycle for grayscale stimuli) or the relatively central location of their flickering stimulus. The high contrast ratio of the OLED, which can show true blacks, may have also contributed to some residual visibility in our study. Given that we obtained a robust RIFT signal for central stimuli, there is likely some room to reduce the contrast of the flickering disc to minimize visibility. In future work, we plan to systematically investigate the factors modulating RIFT (in)visibility with OLEDs.

In sum, in the current pilot study, we provide a working recipe for EEG-based RIFT using affordable hardware. We hope these findings open the door to broader applications of RIFT in basic research as well as with BCIs (Brickwedde et al., 2022; see also Ladouce et al., 2022) or clinical applications, where cost and portability matter. Open questions remain around the precise retinotopic tuning of RIFT responses in EEG, the limits of flicker (in)visibility on OLEDs, and the replicability of attention-related modulations in this setup.

## Footnotes

1 For the first three participants, as a manipulation check, we instead presented a central binary (on/off) SSVEP flicker at 30 Hz (results not shown).

2 Another critical system optimization addressed an obscure Ubuntu 24.04 bug that prevents proper page-flipping unless the top–right Ubuntu notification area is clicked, this workaround then lasts until the next login (Kleiner, 2024). Without this fix, occasional multi-frame drops (of 4–5 frames) caused visible flashes.

